# Genotype 1 Japanese encephalitis virus predominates in nature in China

**DOI:** 10.1101/591339

**Authors:** Nan Shao, Shihong Fu, Fan Li, Weijia Zhang, Qianying Wang, Wenwen Lei, Ying He, Yuxi Cao, Guodong Liang, Huanyu Wang

## Abstract

**Introduction:** Japanese encephalitis virus (JEV), in the genus *Flavivirus*, family *Flaviviridae*, is the leading cause of viral encephalitis in the Asian-Pacific region. JEV has a wide range of vector hosts, including mosquitoes, swine, and wading birds. *Culex* species, especially *Culex tritaeniorhynchus*, are the main vectors for JEV transmission. JEV has been classified into five genotypes (G1–G5). JEV G1, G3, and G5 isolates have been found in China.

**Methods:** More than 200,000 mosquitoes collected in 14 provinces in China from 2004 to 2016 were examined for JEV using a TaqMan real-time RT-PCR assay. The JEV envelope (E) gene was sequenced from positive pools. The nucleic acid sequences were analyzed and aligned using ClustalX ver. 2.0. Representative JEV envelope (E) gene sequences were downloaded from GenBank and compared with the newly obtained JEV sequences. MEGA 5.05 was used to generate a phylogenetic tree based on the JEV sequences.

**Results:** More than 200,000 mosquitoes were divided into 3107 pools to test for JEV. Of the pools, 9.04% (281/3107) were positive. The JEV obtained from mosquitoes included three genotypes (G1, G3, and G5). JEV G1 was detected from different provinces during 2004–2016. G1 was the dominant genotype circulating in nature, comprising 98.93% (278/281). JEV G3 was detected in Yunnan and Chongqing Provinces before 2010; JEV G5 was detected only in Tibet. No pools were positive for JEV G2 and G4. *C*. *tritaeniorhynchus* was the major mosquito species and the principal vector for transmitting JEV in China. The results of the TaqMan real-time RT-PCR assay and phylogenetic analysis consistently indicated that all 50 newly identified JEV sequences belonged to JEV G1.

**Conclusion:** G1 is the main genotype of JEV circulating in nature in China. Some G3 and G5 were found, but no G2 or G4.

**Author summary:** Japanese encephalitis virus (JEV) is a member of the genus *Flavivirus*, family *Flaviviridae* and is the major pathogen agent of Japanese encephalitis (JE) in the Asian-Pacific region. The fatality rate of JE is 20–30%. JEV consists of five genotypes (G1-G5). The predominant genotype was G3 before the 20^th^century. G1 and G3 co-circulated in China at the end of the 20^t^^h^and early 21^st^century. JEV G1 is the youngest genotype and has gradually been found in many areas worldwide. Using a sensitive TaqMan real-time RT-PCR assay, this study examined 3107 pools of mosquito. The main genotype of Japanese encephalitis virus in China was G1, with some G3 and G5, but no G2 or G4. *C*. *tritaeniorhynchus* was the major mosquito species and the principal vector for transmitting JEV in China. TaqMan real-time RT-PCR assay and phylogenetic analysis consistently indicated that all 50 newly identified JEV sequences belonged to JEV G1.

## Introduction

Japanese encephalitis virus (JEV), a mosquito-borne flavivirus, is maintained in nature in a zoonotic cycle involving mosquitoes (principally *Culex* species), pigs, water birds, and bats [1-2]. Humans are dead-end hosts because when bitten by infected mosquitoes, the viremia is transient and low-level [3-4]. JEV is the causative agent of Japanese encephalitis (JE), which is prevalent in most Asian countries and parts of the Western Pacific region, with two epidemiological models (epidemic or endemic) [5-6]. An estimated three billion people widely distributed in 24 countries live in areas with a risk of JEV transmission [7-8]. JE has a fatality rate of 20–30% and 30–50% of the survivors suffer irreversible neurological or psychiatric sequelae [9-10].

JEV has a single-stranded, positive-sense RNA genome that is approximately 11 kb in length. The viral genome contains three parts: 5′-end and 3′-end untranslated regions and an open reading frame (ORF). The 5′-end untranslated region has a type I cap structure with m7G(5)ppp(5′)ApUp, but there is no poly(A) tail at the 3′-end. The ORF encodes three structural proteins, the capsid (C), membrane (M), and envelope (E) proteins, and seven nonstructural proteins (NS1, NS2A, NS2B, NS3, NS4A, NS4B, and NS5) [11-12].

JEV has been divided into five different genotypes based on the complete E gene [13-14] and there are epidemic differences among the JEV genotypes. G1, G3, and G5 are mainly prevalent and distributed in Asia. G1 and G3 are found in most Asian countries, while G5 is currently found only in Malaysia [15-16], China [17], and Korea [18]. G2 has been isolated in South Korea, Southern Thailand, Malaysia, Indonesia, Singapore, Papua New Guinea, and Northern Australia since 1951, while G4 has been isolated only from Indonesia [16, 19]. The dominant genotype in Asia has gradually shifted from G3 to G1 since the 1990s [20]. The distribution of G1 is widespread and has diffused significantly faster between regions than G3 [21]. During 2002–2004, G1 and G3 co-circulated according to JEV surveillance of mosquitoes in China [22]. Previous studies have shown that JEV G1 was the youngest genotype [20, 23]. In this study, we used a rapid, specific, and sensitive TaqMan real-time RT-PCR assay to detect and differentiate JEV from mosquitoes collected from 14 provinces in China during 2004–2016 to comprehensively examine the distribution of JEV.

## Methods

### Mosquito collection

Mosquitoes were collected from 14 provinces (Xinjiang, Qinghai, Yunnan, Tibet, Sichuan, Chongqing, Shaanxi, Henan, Shanxi, Hunan, Hubei, Shanghai, Shandong, and Liaoning) of China at different sites (all collection was done on public land) from 2004 to 2016. The mosquitoes were pooled by species (up to 100 mosquitoes per pool) for each collection site. The mosquito pools collected in 2016 were added to 1.5 mL of minimum essential medium (MEM; Gibco, Invitrogen, Carlsbad, CA, USA) supplemented with 5% PS (1000 unit/mL penicillin and 100 μg/mL streptomycin; Gibco, Invitrogen), 1% glutamine, and 0.12% NaHCO_3_, and ground using TissueLyser (QIAGEN, Hilden, Germany) [17]. The mosquito homogenates were centrifuged at 14,000 rpm/min for 30 min at 4°C and stored at –80°C for later use. The clarified supernatant of the homogenates of mosquito pools collected between 2004 and 2015 were previously stored at –80°C.

### RNA extraction and virus detection

Viral RNA was extracted from 140 µL of clarified supernatant of the homogenates using the QIAamp® Viral RNA Mini Kit (QIAGEN), according to the manufacturer’s specifications. TaqMan real-time RT-PCR was conducted using AgPath-ID™ One-step RT-PCR Reagents (Applied Biosystems, Foster City, CA, USA) and performed on an Mx3000P QPCR System (Stratagene, La Jolla, CA, USA). Each RT-PCR master mix reaction used 12.5 µL 2× RT-PCR buffer, 1 µL forward and reverse primers (both 10 pmol/µL), 1 µL TaqMan probe (5 pmol/µL), 1 µL 25× RT-PCR Enzyme Mix, and 2 µL RNA template for a total volume of 25 µL. qPCR was performed at 45°C for 10 min, 95°C for 10 min, and 40 cycles of 95°C for 15 s and 60°C for 1 min. The cut off cycle threshold (Ct) value was set at 35. Six sets of primers and probes were used in the TaqMan real-time RT-PCR detection system for detecting JEV, as described previously [24].

The positive extraction control consisted of supernatant from JEV-infected BHK cells. The JEV strains were GZ56 (HM366552), P3 (JEU47032), and XZ0934 (JF91589) for G1, G3, and G5, respectively. And the negative extraction control consisted of supernatant from uninfected BHK cells. Double-distilled H_2_O was used as a no-template control.

### PCR amplification and DNA sequencing

First strand cDNA was synthesized from JEV RNA from JEV-positive pools using the PrimeScript™ One Step RT-PCR Kit ver. 2 (Dye Plus; TaKaRa, Dalian, Japan), according to the manufacturer’s instructions. Three pairs of primers were used to obtain the complete E gene: primers JE-E1-F (5′-AYCCTGGYTAYGCTTTCCT-3′, 868–886) and JE-E1-R (5′-GTTCAGTCCACTCCTTGGYTCACA-3′, 1545–1568) for E1; primers JE-E2-F (5′-TACWGTAACWCCMAATGCTC-3′, 1478–1497) and JE-E2-R (5′-CCAGCACCTTTGAGTTRG-3′, 2074–2091) for E2; and primers JE-E3-F (5′-GCTGGTRACAGTRAACCCCTTCGT-3′, 2036–2059) and JE-E3-R (5′-TTTCTTGTGATGTCAATGGC-3′, 2490–2509) for E3. The amplified products were visualized by agarose gel electrophoresis (1%) and positive PCR products were sent to Sangon Biotech (Shanghai) for sequencing.

### Infection rate and positive pools

The minimum infection rate (MIR) of *C*. *tritaeniorhynchus* was estimated using the following formula: (number of JEV-positive pools / total number of mosquitoes tested) × 1000 (https://www.cdc.gov/westnile/resourcepages/mosqsurvsoft.html).

Positive pools were estimated as follows: (number of JEV-positive pools by TaqMan real-time PCR / total pools of mosquitoes tested) × 100 [25].

### Phylogenetic analysis

To construct a phylogenetic tree, representative JEV sequences were downloaded from GenBank. In addition, the E genes of newly sequenced JEV from mosquitoes in China from 2004 to 2016 were included. All newly generated sequences were assembled and manually adjusted using the SeqMan program of DNASTAR (DNASTAR, Madison, WI, USA). The multiple sequence alignment of E gene sequences was analyzed using ClustalX ver. 2.0 [26]. The maximum likelihood (ML) method in MEGA 5.05 [27] was used to construct phylogenetic trees based on the E gene. Tamura-Nei distances and the gamma distribution with invariant sites (G+I) formed the best nucleotide substitution model as recommended by the models function in MEGA. The robustness of the tree was evaluated using 1,000 bootstrap replications. Murray Valley encephalitis virus (MVEV) strain MVE-1-51 was used as the outgroup.

## Results

### Total positive rate

A total of 209,297 mosquitoes representing five genera were collected from 14 provinces in China during 2004–2016 and formed 3107 pools (Figure 1). Of these, 281 pools (9.04%) from 12 provinces (except Qinghai and Xinjiang) were JEV-positive (Ct < 35).

**Figure 1.**
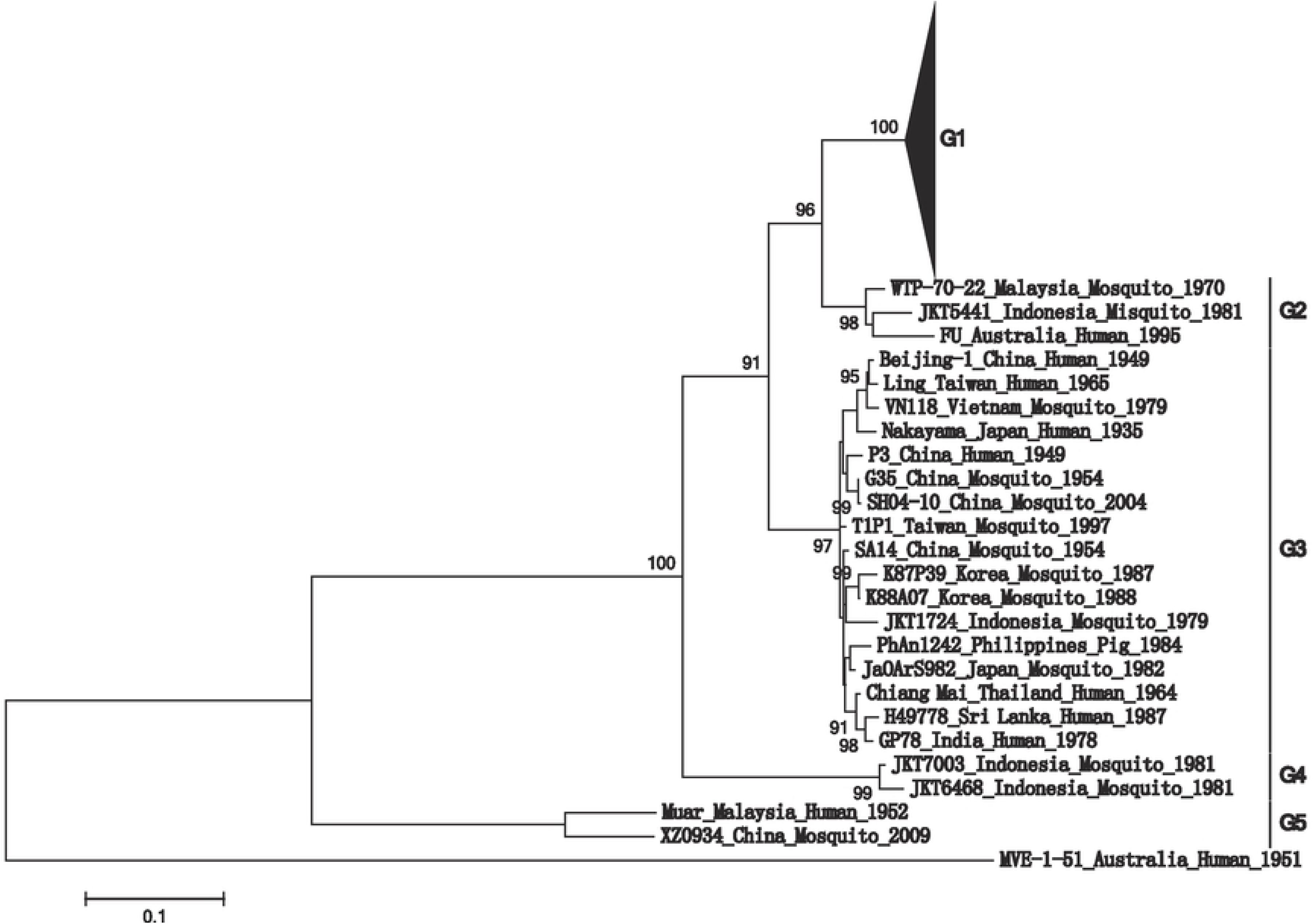
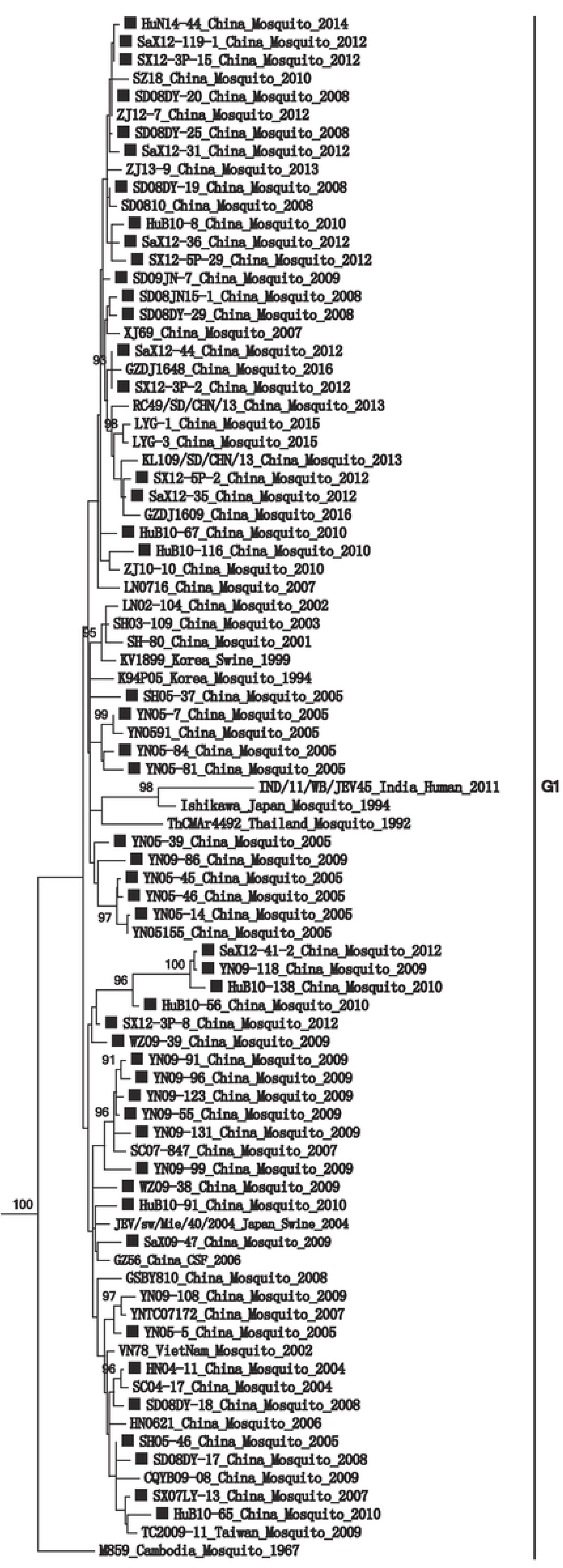
Geographical distribution of Japanese encephalitis virus (JEV) in mainland China. The distribution of JEV was visualized by using the software ArcGIS 9.3(ESRI Inc., Redlands, CA, USA). Abbreviations: AH, Anhui; BJ, Beijing; CQ, Chongqing; FJ, Fujian; GD, Guangdong; GS, Gansu; GZ, Guizhou; GX, Guangxi; HuN, Hunan; HuB, Hubei; HN, Henan; HB, Hebei; HaiN, Hainan; HLJ, Heilongjiang; IMG, Inner Mongolia; JL, Jilin; JS, Jiangsu; LN, Liaoning; NX, Ningxia; QH, Qinghai; SC, Sichuan; SaX, Shaanxi; SX, Shanxi; SD, Shandong; SH, Shanghai; TJ, Tianjin; XJ, Xinjiang; XZ, Tibet; YN, Yunnan; ZJ, Zhejiang

### Distribution of JEV genotypes in mosquitoes

The JEV obtained from the mosquitoes in China included three genotypes (G1, G3, and G5). The distribution of G1 has expanded continuously. JEV G1 was detected in Sichuan and Henan provinces in 2004, followed by Shanghai and Yunnan in 2005, and subsequently in other provinces. Moreover, of the 281 positive pools, 278 (98.9%) pools were JEV G1. JEV G1 was detected in Yunnan, Sichuan, Chongqing, Shaanxi, Henan, Hunan, Hubei, Shanxi, Shandong, Liaoning, and Shanghai Provinces and Tibet. The proportions of positive pools in Henan in 2004, Yunnan in 2005, and Shandong in 2008 were above 30%, those for Shanghai in 2005, Hubei in 2010, Shanxi in 2012, and Shandong in 2016 were above 20%, and those from Sichuan in 2004, Yunnan in 2009, and Shaanxi in 2012 were above 15%; the others were less than 10% (Table 1).

The results indicated that the infection rate of mosquitoes in China by JEV G1 has been high since 2004. JEV G3 was detected from mosquitoes collected in Yunnan in 2005 and Chongqing in 2008. However, no JEV G3 has been detected from mosquitoes collected in these two regions or any other area since then. Moreover, JEV G5 was detected only in Tibet in 2009. This pool positive for G5 was the same pools from which the XZ0934 strain was isolated from Tibet in 2009. Although JEV is widely distributed in nature in China, no JEV was detected in Qinghai or Xinjiang.

### Infection rate of C. tritaeniorhynchus

More than 200,000 mosquitoes were collected, of which 44.71% were *C*. *tritaeniorhynchus*. Of the JEV-positive pools, 82.56% (232/281) were *C*. *tritaeniorhynchus*, indicating that *C*. *tritaeniorhynchus* is the major vector for JEV transmission. No *C*. *tritaeniorhynchus* was collected in Xinjiang and Qinghai. The MIR of *C*. *tritaeniorhynchus* ranged from 0.41 to 9.99 in different provinces and the highest MIR value was found in Yunnan in 2005 (Table 2).

### Phylogenetic analysis of JEV

The E fragments of all positive specimens were amplified by qPCR and 63 JEV G1 sequences were obtained. Thirteen sequences were excluded because they were found in the same pools in previous studies. The sequence alignment showed that these 13 sequences were identical to the published sequences. Therefore, 50 new E gene sequences of JEV G1 were obtained.

A ML tree was constructed based on the complete E gene to establish the phylogenetic relationships of JEV (G1–G5) strains downloaded from GenBank and the 50 new JEV sequences. MVEV strain MVE-1-51 was used as the outgroup. Figure 2 shows the phylogenetic tree constructed from the data in Table 3. The new JEV sequences were dispersed in the two main G1 lineages. No obvious geographical aggregation was observed for the JEV strains from China.

**Figure 2.**
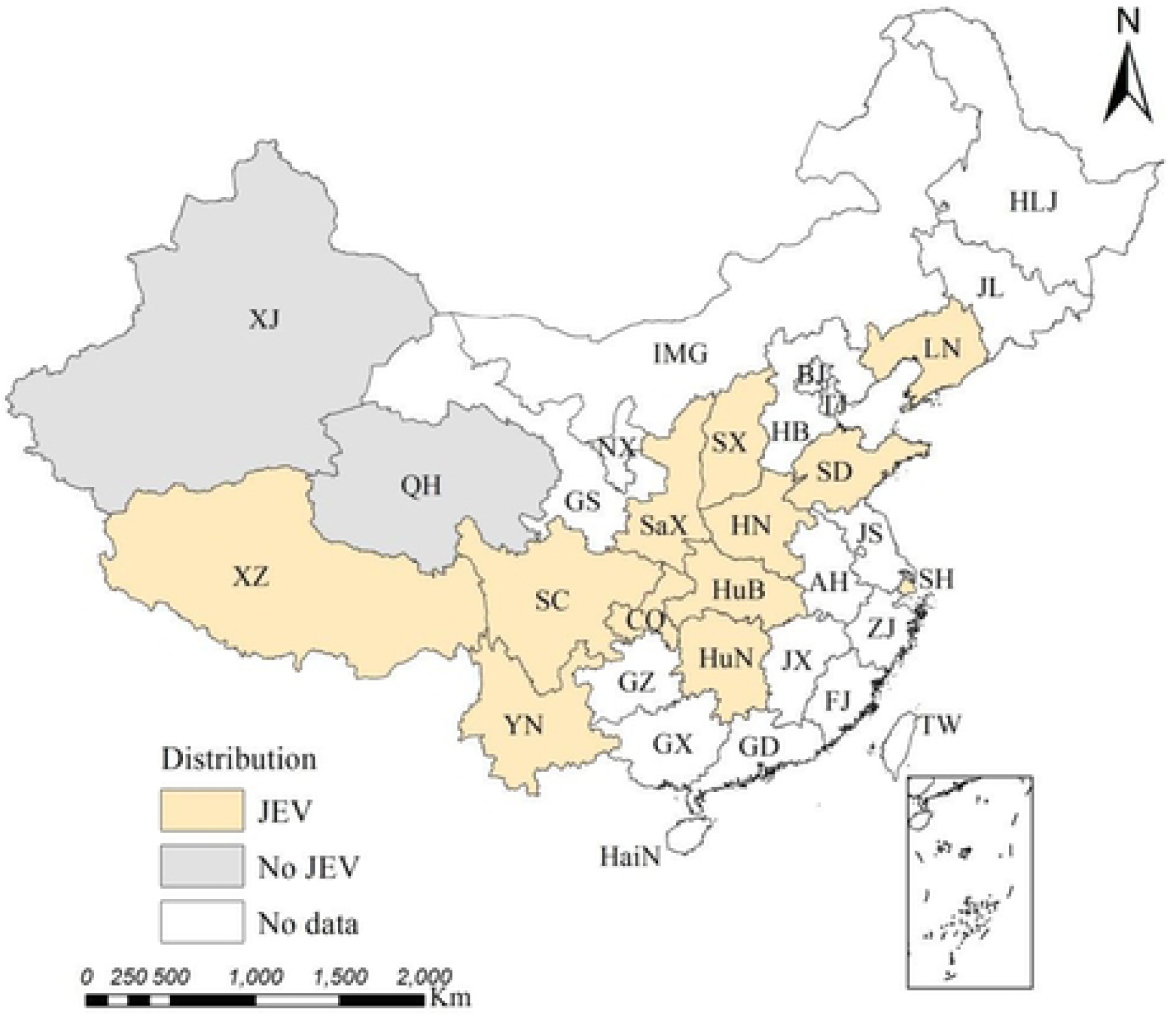
Phylogenetic analysis based on the envelope (E) gene of JEV. The newly sequenced JEV sequences used in this study are marked with squares. The English in this document has been checked by at least two professional editors, both native speakers of English. For a certificate, please see: http://www.textcheck.com/certificate/l0Th6G

## Discussion

The prevalence and distribution of JEV in nature in China involved G1, G3, and G5. No G2 or G4 was found. All 50 JEV sequences newly identified in this study belonged to G1 based on the phylogenetic analysis and TaqMan real-time RT-PCR assay. Consistent with previous studies, *C*. *tritaeniorhynchus* was the most important JEV-transmitting mosquito, which transmits the virus in a mosquito–pig cycle and human infection [28]. No JEV was detected in Qinghai and Xinjiang Provinces.

Historically, cell cultures and traditional PCR are used to verify JEV. In this study, we not only detected JEV but also distinguished the genotypes based on the established TaqMan real-time RT-PCR assay. This method showed high specificity, sensitivity, and reproducibility, and reflected the distribution of JEV reliably and accurately in mainland China. In addition, two target fragment sequences of G3 were not obtained due to their high Ct values with qPCR amplification.

Although the mosquitoes in this study were found in a wide range of geographical areas in China over a long period, all of the obtained information was limited to mosquitoes. Therefore, we need to obtain additional data from different hosts, such as humans, pigs, and birds. Only comprehensive information can promote further understanding of the distribution of JEV and changes in its genotypes. Some studies have confirmed G5 in South Korea in recent years [18, 29]. Therefore, the potential impact of JEV G5 from a neighboring country cannot be neglected.

Encephalitis caused by mosquito-borne JEV is serious. JE is classified as a Class B infectious disease in China. Long-term passive surveillance data indicate that cases of JE have been reported throughout mainland China, except Xinjiang, Qinghai, and Tibet [30-31]. At present, there are no effective antiviral drugs against JE and vaccination is regarded as the most economical and effective preventive method. Only four JE vaccines have been approved by the World Health Organization: JEEV and JEEV Pediatric in India, Live-SA14-14-2 in China, and IMOJEV.MD in Thailand. All of these JE vaccines are based on JEV G3 strains. Some *in vitro* and *in vivo* studies have shown that these JE vaccines derived from G3 strains do not provide adequate protection against G5 [32-33] and have reduced neutralizing capacity against G1 [34].

Therefore, preventive strategies applied to disease control caused by JEV are necessary. More detailed studies are needed to address selection pressure, antigenicity, and the pathogenicity features of JEV following large-scale application of inactivated or live attenuated vaccines derived from G3 strains. From a public health perspective, it will also be important to prepare JEV vaccines against G1 and G3 simultaneously or even against the potential threat from JEV G5 and emerging new genotypes.

## Author contribution

Huanyu Wang designed the study; Nan Shao, Shihong Fu, Fan Li, Weijia Zhang and Qianying Wang performed the experiments; Nan Shao, Shihong Fu, Wenwen Lei and Ying He performed mosquito collection; Yuxi Cao and Guodong Liang contributed to drafting and editing the paper; all authors reviewed the manuscript.

## Acknowledgments

We are grateful to the people from the relevant Centers for Disease Control for participating in sample collection.

## References

[1] Solomon T. Flavivirus encephalitis. N Engl J Med. 2004;351: 370–378.

[2] Wang JL, Pan XL, Zhang HL, Fu SH, Wang HY, Tang Q, et al. Japanese Encephalitis Viruses from Bats in Yunnan, China. Emerg Infect Dis. 2009;15: 939–942.

[3] Solomon T. Control of Japanese encephalitis--within our grasp?. N Engl J Med. 2006;355: 869–871.

[4] Seo H, Kim HC, Klein TA, Ramey AM, Lee J, Kyung S, et al. Molecular Detection and Genotyping of Japanese Encephalitis Virus in Mosquitoes during a 2010 Outbreak in the Republic of Korea. PLOS One. 2013;8: e55165.

[5] Wang HY, Liang GD. Epidemiology of Japanese encephalitis: past, present, and future prospects. Ther Clin Risk Manag. 2015;11: 435–448.

[6] Erlanger T E, Weiss S, Keiser J, Utzinger J, Wiedenmayer K. Past, Present, and Future of Japanese Encephalitis. Emerg Infect Dis. 2009;15: 1–7.

[7] Heffelfinger JD, Li X, Batmunkh N, Grabovac V, Diorditsa S, Liyanage JB, Pattamadilok S, et al. Japanese Encephalitis Surveillance and Immunization - Asia and Western Pacific Regions, 2016. MMWR. Morb Mortal Wkly Rep. 2017;66: 579–583.

[8] Campbell GL, Hills SL, Fischer M, Jacobson JA, Hoke CH, Hombach JM, et al. Estimated global incidence of Japanese encephalitis: a systematic review. Bull World Health Organ. 2011;89: 766-774, 774A.

[9] Tom Solomon, Dung NM, Rachel Kneen, Mary Gainsborough, David W Vaughn, Khanh VT. Japanese encephalitis. J Neurol Neurosurg Psychiatry. 2000;68: 405–415.

[10] Zhang JS, Zhao QM, Guo XF, Zuo SQ, Cheng JX, Jia N, et al. Isolation and genetic characteristics of human genotype 1 Japanese encephalitis virus, China, 2009. PLOS One. 2011;6: e16418.

[11] Sumiyoshi H, Mori C, Fuke I, Morita K, Kuhara S, Kondou J, et al. Complete nucleotide sequence of the Japanese encephalitis virus genome RNA. Virology. 1987;161: 497–510.

[12] Chambers TJ, Hahn CS, Galler R, Rice CM. Flavivirus genome organization, expression, and replication. Annu Rev Microbiol. 1990;44: 649–688.

[13] Solomon T, Ni H, Beasley DW, Ekkelenkamp M, Cardosa MJ, Barrett AD. Origin and evolution of Japanese encephalitis virus in southeast Asia. J Virol. 2003;77: 3091–3098.

[14] Uchil P D, Satchidanandam V. Phylogenetic analysis of Japanese encephalitis virus: envelope gene based analysis reveals a fifth genotype, geographic clustering, and multiple introductions of the virus into the Indian subcontinent. Am J Trop Med Hyg. 2001;65: 242–251.

[15] Hale JH, Lim KA, Chee PH. Japanese type B encephalitis in Malaya. Ann Trop Med Parasitol. 1952;46: 220–226.

[16] Mohammed M A F, Galbraith SE, Radford AD, Dove W, Takasaki T, Kurane I, et al. Molecular phylogenetic and evolutionary analyses of Muar strain of Japanese encephalitis virus reveal it is the missing fifth genotype. Infect Genet Evol. 2011;11: 855–862.

[17] Li MH, Fu SH, Chen WX, Wang HY, Guo YH, Liu QY, et al. Genotype v Japanese encephalitis virus is emerging. PLOS Negl Trop Dis. 2011;5: e1231.

[18] Takhampunya R, Kim H, Tippayachai B, Kengluecha A, Klein T A, Lee W, et al. Emergence of Japanese encephalitis virus genotype V in the Republic of Korea. Virol J. 2011;8: 449.

[19] Schuh AJ, Ward MJ, Brown A J L, Barrett A D T. Phylogeography of Japanese encephalitis virus: genotype is associated with climate. PLOS Negl Trop Dis. 2013;7: e2411.

[20] Pan XL, Liu H, Wang HY, Fu SH, Liu HZ, Zhang HL, et al. Emergence of Genotype I of Japanese Encephalitis Virus as the Dominant Genotype in Asia. J Virol. 2011;85: 9847–9853.

[21] Pham TT, Meng S, Sun Y, Lv W, Bahl J. Inference of Japanese encephalitis virus ecological and evolutionary dynamics from passive and active virus surveillance. Virus Evol. 2016;2: w9.

[22] Wang HY, Takasaki T, Fu SH, Sun XH, Zhang HL, Wang ZX, et al. Molecular epidemiological analysis of Japanese encephalitis virus in China. J Gen Virol. 2007;88: 885–894.

[23] Gao X, Liu H, Li M, Fu S, Liang G. Insights into the evolutionary history of Japanese encephalitis virus (JEV) based on whole-genome sequences comprising the five genotypes. Virol J. 2015;12: 43.

[24] Shao N, Li F, Nie K, Fu SH, Zhang WJ, He Y, et al. TaqMan Real-time RT-PCR Assay for Detecting and Differentiating Japanese Encephalitis Virus. Biomed Environ Sci, 2018;31: 208–214.

[25] Chu H, Wu Z, Chen H, Li C, Guo X, Liu R, et al. Japanese Encephalitis Virus Infection Rate and Detection of Genotype I From Culex tritaeniorhynchus Collected From Jiangsu, China. Vector Borne Zoonotic Dis. 2017;17:503–509.

[26] Larkin MA, Blackshields G, Brown NP, Chenna R, Mcgettigan PA, Mcwilliam H, et al. Clustal W and Clustal X version 2.0. Bioinformatics. 2007;23: 2947–2948.

[27] Tamura K, Peterson D, Peterson N, Stecher G, Nei M, Kumar S. MEGA5: Molecular Evolutionary Genetics Analysis Using Maximum Likelihood, Evolutionary Distance, and Maximum Parsimony Methods. Mol Biol Evol. 2011;28: 2731–2739.

[28] Liang G, Li X, Gao X, Fu S, Wang H, Li M, et al. Arboviruses and their related infections in China: A comprehensive field and laboratory investigation over the last 3 decades. Rev Med Virol. 2018;28: e1959.

[29] Kim H, Cha G, Jeong YE, Lee W, Chang KS, Roh JY, et al. Detection of Japanese Encephalitis Virus Genotype V in Culex orientalis and Culex pipiens (Diptera: Culicidae) in Korea. PLOS One. 2015;10: e116547.

[30] Wang H, Li Y, Liang X, Liang G. Japanese encephalitis in mainland china. Jpn J Infect Dis. 2009;62: 331–336.

[31] Zheng Y, Li M, Wang H, Liang G.Japanese encephalitis and Japanese encephalitis virus in mainland China. Rev Med Virol. 2012;22: 301–322.

[32] Cao L, Fu S, Gao X, Li M, Cui S, Li X, et al. Low Protective Efficacy of the Current Japanese Encephalitis Vaccine against the Emerging Genotype 5 Japanese Encephalitis Virus. PLOS Negl Trop Dis. 2016;10: e4686.

[33] Moi ML, Yagasaki K, Tajima S, Nakayama E, Takasaki T, Saijo M, et al. In vitro growth, pathogenicity and serological characteristics of the Japanese encephalitis virus genotype V Muar strain. J Gen Virol. 2015;96: 2661–2669.

[34] Fan YC, Chen JM, Chiu HC, Chen YY, Lin JW, Shih CC, et al. Partially neutralizing potency against emerging genotype I virus among children received formalin-inactivated Japanese encephalitis virus vaccine. PLOS Negl Trop Dis. 2012;6: e1834.

